# Structural basis for the peptidoglycan editing activity of YfiH

**DOI:** 10.1101/2021.12.07.471703

**Authors:** Meng-Sheng Lee, Kan-Yen Hsieh, Chiao-I Kuo, Szu-Hui Lee, Chung-I Chang

## Abstract

Bacterial cells are encased in peptidoglycan (PG), a polymer of disaccharide N-acetyl-glucosamine (GlcNAc) and N-acetyl-muramic acid (MurNAc) cross-linked by peptide stems. PG is synthesized in the cytoplasm as UDP-MurNAc-peptide precursors, of which the amino-acid composition of the peptide is unique, with L-Ala added at the first position in most bacteria but L-Ser or Gly in some bacteria. YfiH is a PG-editing factor whose absence causes misincorporation of L-Ser instead of L-Ala into peptide stems; but its mechanistic function is unknown. Here we report the crystal structures of substrate-bound and product-bound YfiH, showing that YfiH is a cytoplasmic amidase that controls the incorporation of the correct amino acid to the nucleotide precursors by preferentially cleaving the nucleotide precursor byproduct UDP-MurNAc-L-Ser. This work reveals an editing mechanism in the cytoplasmic steps of peptidoglycan biosynthesis.

## Introduction

Peptidoglycan (PG) is a mesh-like polymer of sugars and amino acids uniquely formed in the extracytoplasmic space of bacterial cells. The exoskeleton-like structure of PG maintains cell shape and protects the cells against high turgor pressure (Cabeen and Jacobs-Wagner 2005). The polymeric PG consists of linear sugar chains of alternating N-acetylglucosamine (GlcNAc) and N-acetylmuramic acid (MurNAc), connected by a β-(1,4)-glycosidic bond. Each MurNAc is attached with a tetrapeptide or a pentapeptide stem. In *E. coli*, the stem peptide of PG contains L-alanine, D-glutamic acid, *meso*-diaminopimelic acid (*m*-DAP), and D-alanine. Cross-linking between the stem tetrapeptides, in the case of *E. coli*, of different sugar chains contributes to the 3D mesh-like structure formation; the crosslinks are normally formed between the D-alanine of one tetrapeptide and the *m*-DAP of another tetrapeptide by a peptide bond (Vollmer et al. 2008).

Biosynthesis of PG is initiated in the cytoplasm of bacterial cells. The cytoplasmic steps of PG biosynthesis consist of five sets of reactions, which lead to the formations of (I) UDP-GlcNAc from fructose-6-phosphate, (II) UDP-MurNAc from UDP-GlcNAc, (III) D-Glu from L-Glu, (IV) dipeptide D-Ala-D-Ala from L-Ala, and (V) UDP-MurNAc-L-Ala-D-Glu-*m-*DAP-D-Ala-D-Ala (UDP-MurNAc-pentapeptide) from UDP-MurNAc (Barreteau et al. 2008). UDP-MurNAc-pentapeptide is assembled by sequential ligation of L-Ala, D-Glu, *m*-DAP, and D-Ala-D-Ala to UDP-MurNAc, which are ATP-dependent reactions catalyzed by four amino-acid ligases MurC, MurD, MurE, and MurF, respectively (Barreteau et al. 2008). Once synthesized, UDP-MurNAc-pentapeptide is then transferred to the inner side of cytoplasmic membrane to form lipid-anchored intermediates, of which the final product Lipid II is transported to the outer side of the cytoplasmic membrane to be used by the PG synthases and transpeptidases for polymerization and crosslinking.

The specificities of the Mur ligases are not absolute; for example, although the preferred substrate of MurC is L-Ala, L-Ser or Gly can be added with lower efficiencies *in vitro* (Liger et al. 1995; Emanuele et al. 1996). It has been shown that the MurC ligases isolated from two bacterial species with different amino acids at the first position of PG peptide stem exhibit similar substrate specificities (Mahapatra et al. 2000). A previous work has shown that YfiH (also known as PgeF) is a PG-editing factor with a role in maintaining specific PG composition in *E. coli (Parveen and Reddy 2017)*. Absence of *yfiH* leads to incorporation of L-Ser into the first position of the stem peptide, which is normally occupied by L-Ala, resulting in β-lactam sensitivity, altered cell morphology, and reduced PG synthesis (Parveen and Reddy 2017). Here we elucidate the molecular mechanism of YfiH. We report the crystal structures of YfiH bound to a trapped endogenous UDP-MurNAc, as well as to the preferred substrate UDP-MurNAc-L-Ser. Our results show that YfiH forms an extended L-shaped binding groove for UDP-MurNAc-monopeptide and is a cytoplasmic amidase specific for hydrolyzing UDP-MurNAc-L-Ser, a non-canonical reaction byproduct from the MurC reaction. This work suggests that bacteria possess a cytoplasmic precursor-editing mechanism to maintain specific amino-acid composition of PG.

## Results

### Overall structure of YfiH bound to UDP-MurNAc

We expressed catalytically inactive recombinant *E. coli* YfiH-C107A protein for an original purpose of screening potential substrates or ligands by binding assays and co-crystallography. Purified YfiH-C107A forms highly diffracting crystals; the best crystal diffracted to 1.47 Å resolution, in the space group *P*2_1_2_1_2_1_ with four molecules (chains A-D) per asymmetric unit (Table S1). Chains A and C contain almost all of the YfiH amino-acid residues 2∼243 except for Met1; chains B and D contain residues 3∼243. In chains C and D, residues 81-84 and 79-85, respectively, from a solvent-exposed loop are disordered. The presence of the N-terminal region in the structure indicates that YfiH does not possess a signal peptide and therefore is a cytoplasmic protein. Serendipitously, the electron-density map contained a prominent blob of well-resolved density for an endogenous compound, presumably co-purified with YfiH-C107A and trapped during crystallogenesis. This compound was unambiguously identified as UDP-MurNAc based on the high-resolution map (Fig. 1A). The description and presentation of the complex structure hereafter is based on chain A unless mentioned otherwise.

**Figure 1.**
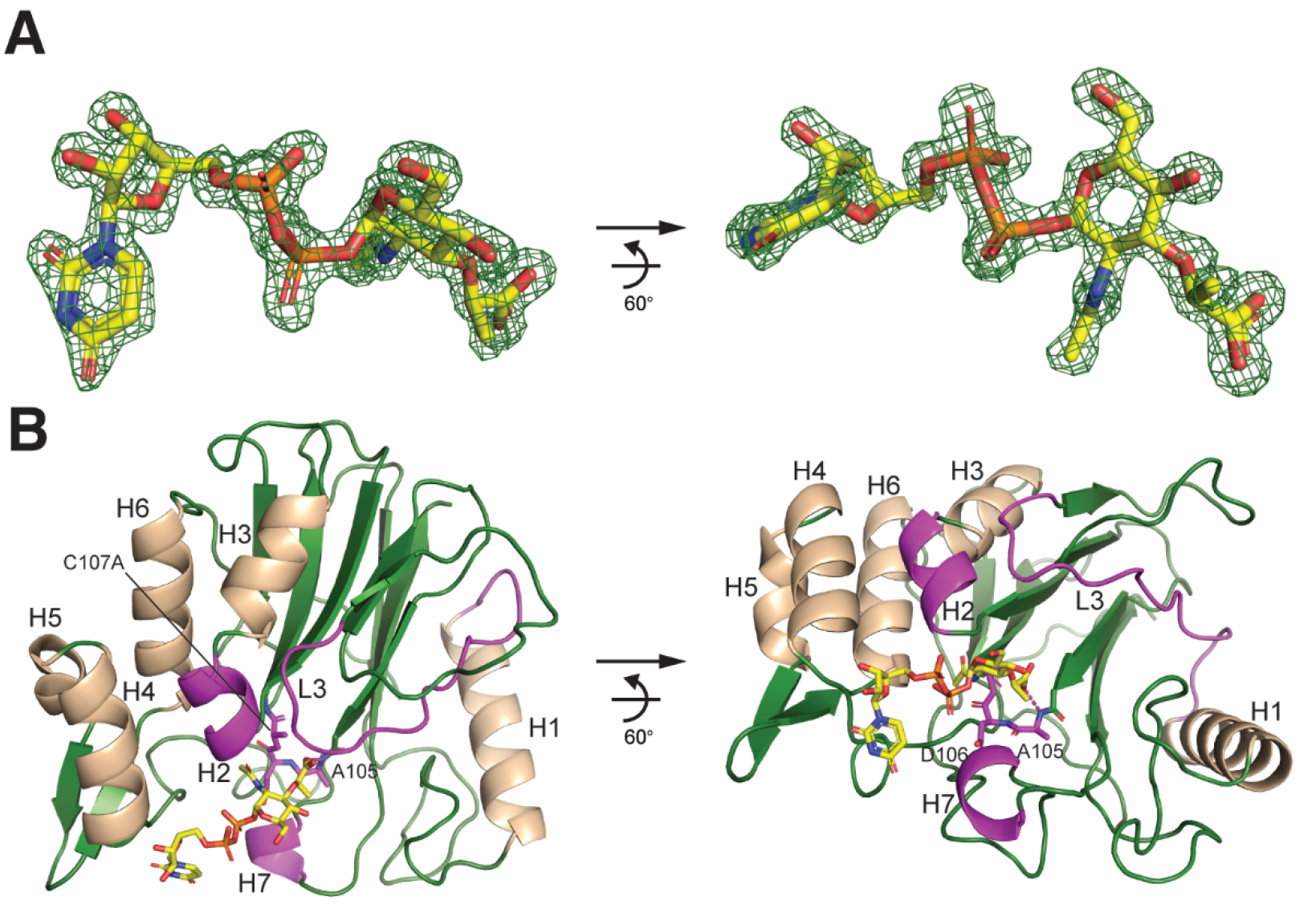
Structure of UDP-MurNAc bound to YfiH-C107A. A. *Fo-Fc* difference Fourier omit map of UDP-MurNAc, shown in sticks and in two views, at 1.47 Å resolution displayed in green isomesh at 4.0 σ level. B. Ribbon diagram of the structure of YfiH, shown in forest green and in two views, with bound UDP-MurNAc in sticks. Helices are colored in wheat. The two helices and the loop interacting with the compound are colored in magenta. The catalytic β-turn is shown in magenta sticks.

The structure of UDP-MurNAc bound *E. coli* YfiH adopts an overall globular structure that is featured by a pair of two central β-sheets sandwiched by multiple α-helices (Figs. 1B and S1), which is similar to that of *Shigella flexneri* YfiH and a bacterial ortholog YlmD (with an r.m.s.d. value of 0.9 Å and 1.3 Å, respectively) (Kim et al. 2006; Cader et al. 2020). UDP-MurNAc is bound to a preformed groove located on one side of the double β-sheets and at a distinct β-turn (a.a. 104-108). The β-turn bridges the two β-sheets; importantly, it harbors the catalytic Cys107 and forms the oxyanion hole by the backbone amides of Ala105 and Asp106 (Fig. 1B). The groove is demarcated by two short helices H2 and H7 and an extensive loop L3 (Fig. EV1), which together guard the catalytic β-turn.

### Interaction of YfiH with UDP-MurNAc-monopeptide

UDP-MurNAc forms a largely extended conformation in the elongated L-shaped binding groove of YfiH, where the uridine moiety forms a sharp angle with the diphosphate linked MurNAc (Fig. 2A). The aromatic ring of uracil packs against the side chain of Arg228 from H7 forming the turning point of the groove. Notably, Arg228 interacts not only with the uracil ring, but also the ribose ring oxygen and the diphosphate of UDP. The UDP diphosphate also makes electrostatic contacts with Trp127 and Arg128 from H2. The sugar ring of MurNAc is stacked with Tyr227 from H7. The lactyl oxygens of MurNAc interact with His71 from L3 and His124, which forms a catalytic triad with Asp89 and Cys107.

**Figure 2.**
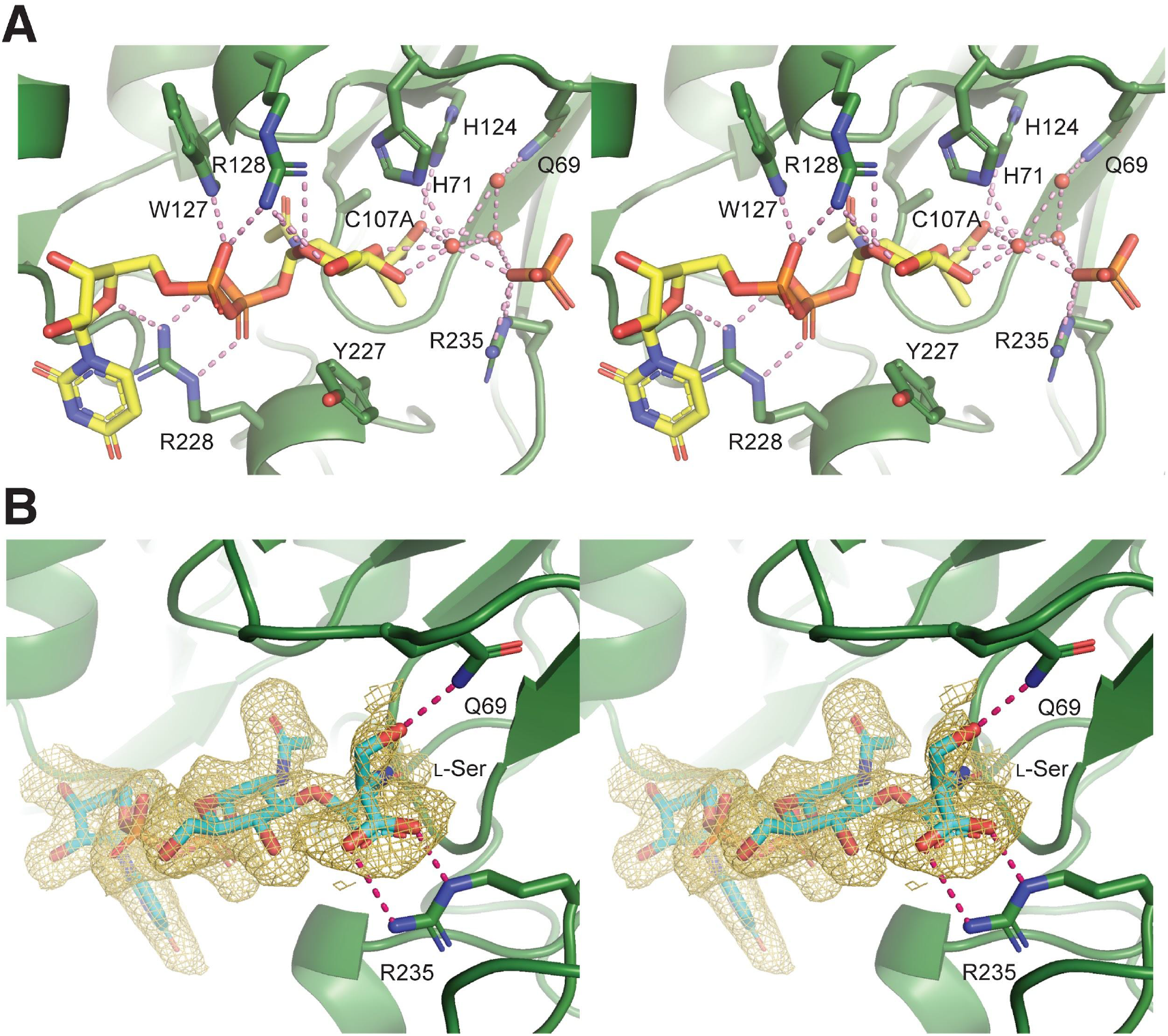
Interactions of UDP-MurNAc and UDP-MurNAc-L-Ser with the binding groove in YfiH. A. Stereoview of the bound UDP-MurNAc and the phosphate ion in the complex structure with YfiH-C107A. Water oxygen atoms are shown in spheres. B. The *Fo-Fc* difference Fourier omit map of UDP-MurNAc-L-Ser bound to YfiH-C107A is displayed in gold isomesh at 1.5 σ level in stereoview. The compounds and the side chains of the interacting residues are shown in sticks. Hydrogen bonds are shown in dashed bonds.

Interestingly, in two of the four YfiH molecules (chains A and B) in the asymmetric unit, a bound phosphate ion is found ∼4.9 Å away from the lactic acid moiety of MurNAc and connected to the free carboxyl group via a hydrogen-bonding network involving several ordered waters (Fig. 2A). The crystallographic structure of these solvent molecules bound to the MurNAc residue strongly suggests the presence of a binding pocket for an amino acid and two putative interacting residues Gln69 and Arg235, which coordinate the water and the phosphate ions, respectively. Therefore, the substrate of YfiH may be a UDP-MurNAc-monopeptide. Based on the genetic observation that absence of *yfiH* leads to the misincorporation of L-Ser instead of L-Ala into PG, our crystallographic results suggest that YfiH may prevent the incorporation of L-Ser by specifically hydrolyzing UDP-MurNAc-L-Ser as the preferred substrate.

To test this idea, we obtained the crystals of YfiH bound to UDP-MurNAc-L-Ser, which was synthesized enzymatically (described below), by washing and soaking the YfiH-UDP-MurNAc crystals with an excess amount of the compound. The co-crystal structure, determined at 1.86 Å resolution (Table S1), shows that Gln69 indeed forms a hydrogen bond with the side-chain hydroxyl of L-Ser of the substrate, whose carboxylic group forms a bidentate salt bridge with Arg235 (Fig. 2B).

### UDP-MurNAc-L-Ser is the preferred substrate of YfiH

We sought to test whether YfiH specifically hydrolyzes UDP-MurNAc-L-Ser into UDP-MurNAc, as well as the role of Gln69 in recognition specificity. To this end, we synthesized UDP-MurNAc-L-Ala (UMA) and UDP-MurNAc-L-Ser (UMS) using recombinant MurC; based on the purified UDP-MurNAc-monopeptides we further synthesized UDP-MurNAc-L-Ala-D-Glu (UMAE) and UDP-MurNAc-L-Ser-D-Glu (UMSE) with recombinant MurD. Purification and quantitative determination of the compounds were performed using HPLC and verified by mass spectrometry (see Methods). We also purified UDP-MurNAc-L-Ala-D-Glu-*m*-DAP (UM-Tri) using an established method (Kohlrausch and Höltje 1991). As expected, wild-type YfiH hydrolyzes UDP-MurNAc-monopeptide into UDP-MurNAc (Figs. 3A and B); the hydrolytic activity requires the catalytic residue Cys107 (Figs. 3C and D). Moreover, the measured specific activity of YfiH on UMS was more than 10 fold greater than UMA (Fig. 3C and Table 1). Removing the side chain of Arg235 significantly decreased the hydrolytic activity (Figs. 2B and 3C). Interestingly, YfiH also hydrolyzed chemically synthesized MurNAc-L-Ser (Fig. S2A). Furthermore, we found that YfiH also cleaves the UM-dipeptides UMAE and UMSE (Figs. S2B and C). Although with significantly lower specific activities, the substrate preference is nevertheless maintained towards UMSE (Figs. 3C and D). Lastly, we could not detect activity of YfiH on UM-tri (Fig. S2D). Taken together, these results demonstrate that YfiH is a cytoplasmic hydrolase specific for the PG precursor UDP-MurNAc-L-serine.

**Figure 3.**
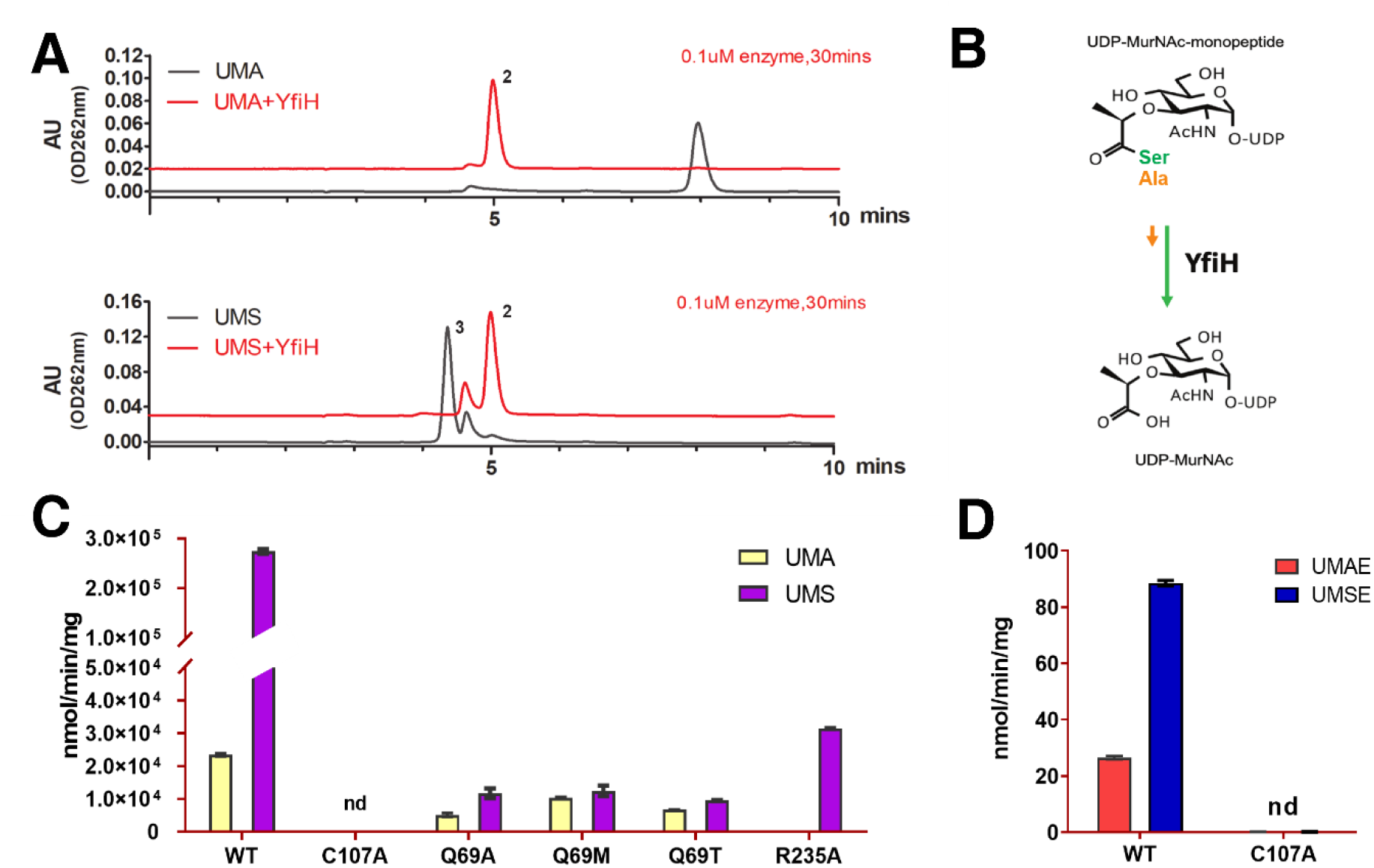
Hydrolyzing activities of YfiH on UDP-MurNAc-peptides and mutational analysis. A. HPLC chromatograms of UDP-MurNAc-L-Ala (UMA; compound 1) and UDP-MurNAc-L-Ser (UMS; compound 3) before and after incubation with YfiH to yield the cleavage product UDP-MurNAc (compound 2). B. Hydrolytic reaction catalyzed by YfiH and the substrate preference for UDP-MurNAc-L-Ser rather than UDP-MurNAc-L-Ala. C. Histogram showing the comparison of the specific activities of wild-type YfiH (YfiH-WT) and mutants on UMA and UMS. D. Histogram showing the comparison of the specific activities of wild-type YfiH and the inactivated mutant C107A on UMAE (UDP-MurNAc-L-Ala-γ-D-Glu), and UMSE (UDP-MurNAc-L-Ser-γ-D-Glu).

**Table 1.**
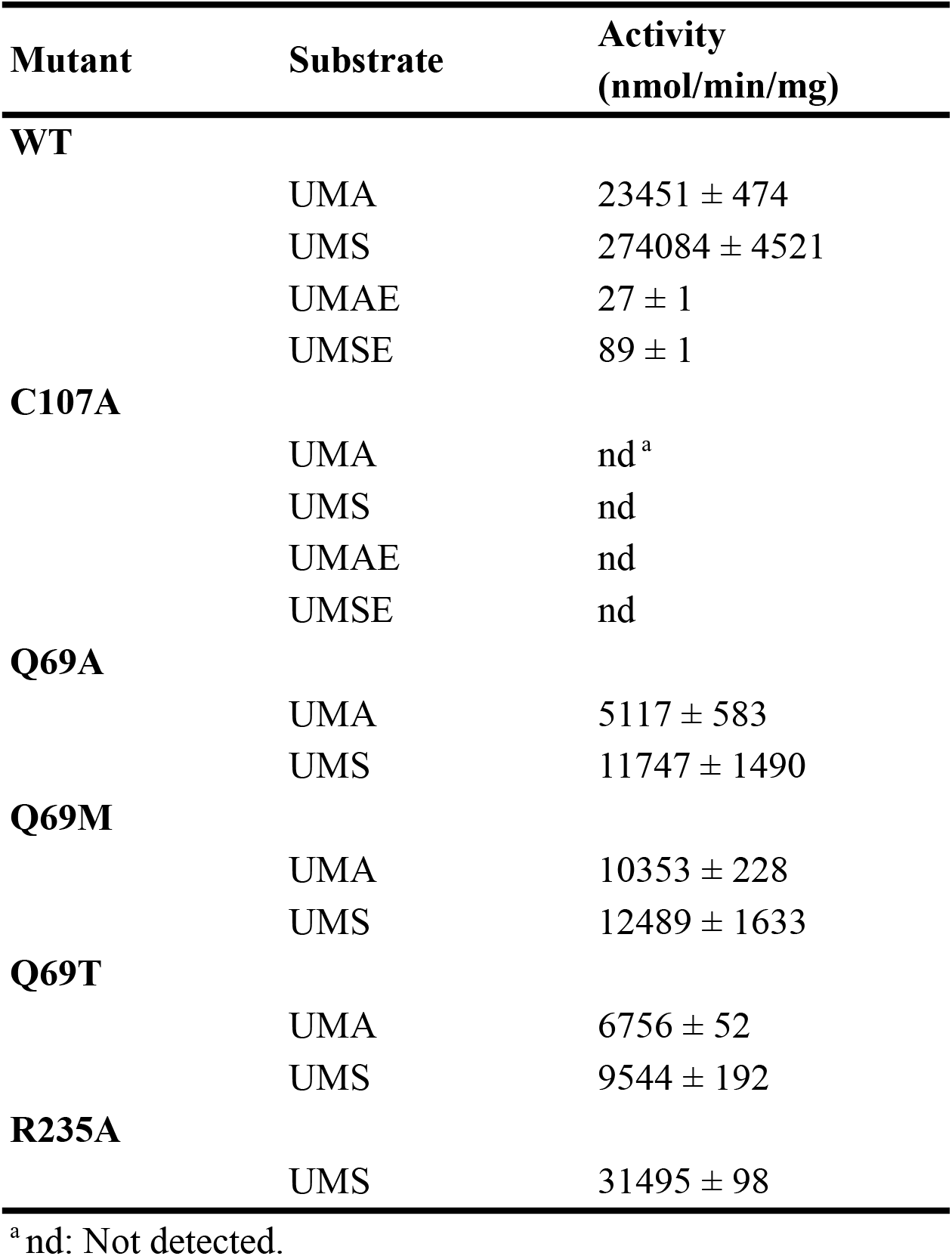
Enzyme activity of YfiH mutants with different UDP-MurNAc derivatives.

We engineered two YfiH mutants Q69A and Q69M to compare their specific activities on the various substrates. The results show that the activity for UMS, but not UMA, was significantly reduced in both Q69A and Q69M mutants whereas the activity for UMA was also affected in Q69A (Figs. 3C and D). Therefore, the long polar side chain of Gln69 may be responsible for interacting with the amino-acid moiety of UDP-MurNAc-monopeptide. Overall, these results demonstrate that Gln69 is the specificity determinant residue.

We have also tested the hydrolytic activity of YfiH on UDP-MurNAc-Gly (UMG) since previous genetic study suggested that YfiH also prevents incorporation of glycine-containing muropeptides (Parveen and Reddy 2017). We attempted to prepare UMG by enzymatic synthesis with MurC, using glycine as a substrate; however, the product yield was very low and purified UMG was extremely unstable in aqueous solution. Therefore, we developed an alternative assay, in which mass spectrometry was used to examine the effect of YfiH on the synthesis of UMG by MurC in the presence of limited amounts of the substrates UM, glycine, and ATP. If YfiH is able to hydrolyze UMG, in the MurC reaction it will prevent accumulation of the product UMG. Concurrently, ATP, which is consumed by the MurC reaction, will be exhausted and eventually no UMG will be present in the reaction. The mass spectra indeed showed that the UMG peak yielded from the MurC reaction disappeared when YfiH, but not YfiH-C107A, was included in the reaction (Figs. 4A-C).

**Figure 4.**
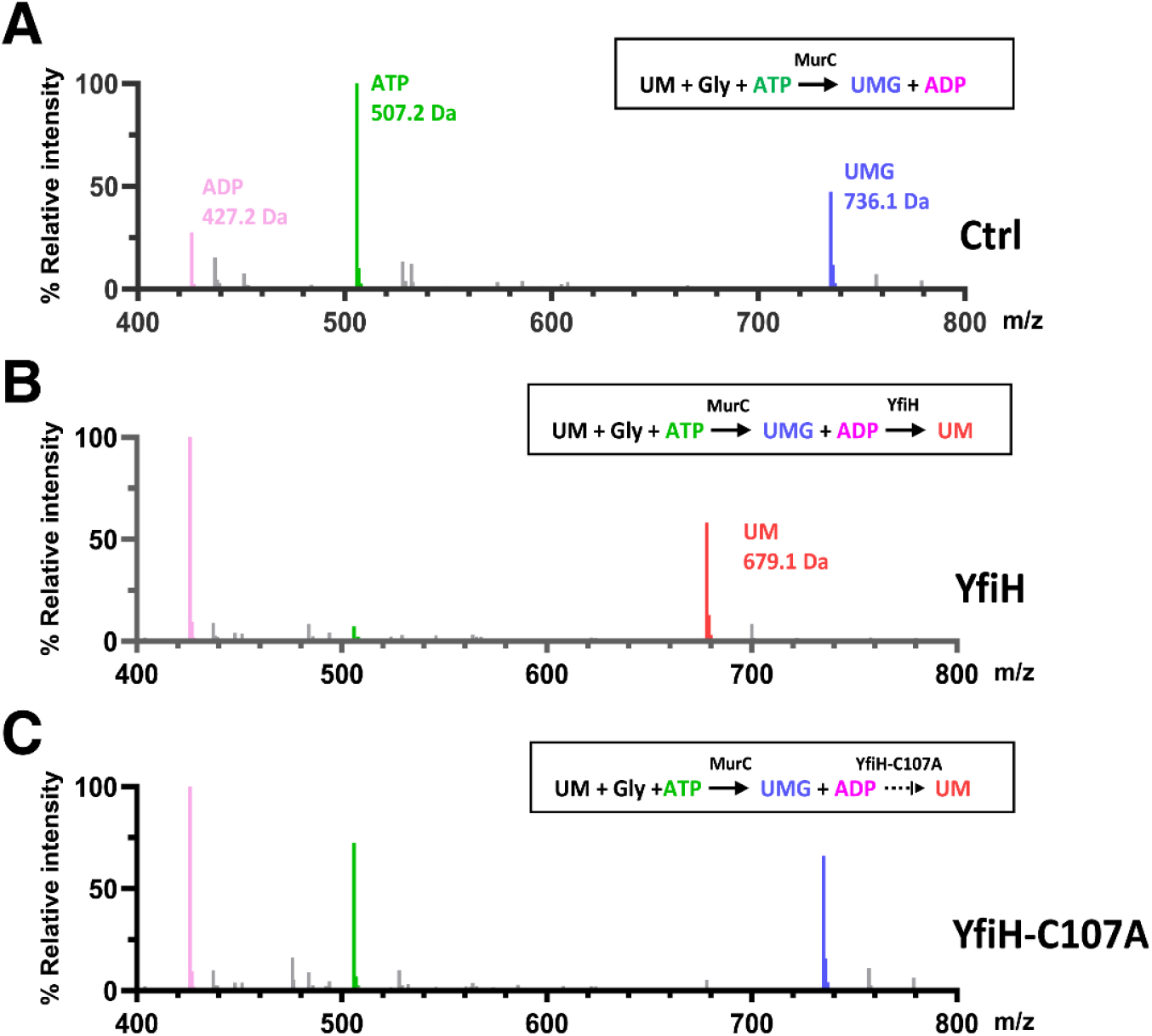
YfiH also hydrolyzes UDP-MurNAc-Gly. A–C. Mass spectra of UDP-MurNAc-Gly (UMG; 736.1 Da) after 24-hour reaction of 1 mM UDP-MurNAc (UM; 679.1 Da), 5 mM ATP, and 18 mM Gly with 20 μM MurC alone (A), co-incubating with 20 μM MurC and 10 μM wild-type YfiH (B), or with 20 μM MurC and 10 μM YfiH-C107A (C).

## Discussion

In this work we show that YfiH is involved in a previously unknown cytoplasmic editing mechanism for biosynthesis of PG (Fig. 5). We show that YfiH hydrolyzes UDP-MurNAc-monopeptides, which are synthesized by MurC, into UDP-MurNAc and amino acids. The preferred substrate of YfiH is UMS, against which the specificity activity of YfiH is more than 10-fold higher than UMA. Although we were unable to determine the specific activity for UMG due to its poor stability, it is likely that YfiH hydrolyzes UMG at a rate comparable, if not inferior, to UMA due to lack of side chain for Gly. The hydrolytic reactions catalyzed by YfiH ensure the incorporation of the specific amino acid L-Ala into the cytoplasmic UDP-precursors of PG. L-Ala is the first amino acid of the peptide moiety of PG in nearly all eubacteria (Schleifer and Kandler 1972). In some bacterial species, however, L-Ser or Gly are added in this position instead of L-Ala (Schleifer and Kandler 1972). In the stepwise assembly of monomeric precursors of PG, formation of the amide bond between UDP-MurNAc and the first amino acid is catalyzed by the MurC ligase (Barreteau et al. 2008). Interestingly, purified MurC ligases from bacteria with the first amino acid of PG peptide being L-Ala or Gly all exhibit preferred catalysis for the synthesis of UDP-MurNAc-L-Ala over UDP-MurNAc-Gly (Liger et al. 1995; Mahapatra et al. 2000). The *E. coli* MurC can take L-Ala, L-Ser, or Gly as substrates; the *K*_*m*_ values for L-Ala, L-Ser, and Gly were 20, 850, and 2500 μM, respectively (Liger et al. 1995). Intriguingly, the MurC ligases from *Mycobacterium leprae* and *Mycobacterium tuberculosis*, which contain Gly and L-Ala in the first position of the stem peptide, respectively, showed similar *K*_*m*_ and *V*_max_ for L-alanine and glycine (Mahapatra et al. 2000). These results suggest that the presence of the species-specific first amino acid in the stem peptide of PG is achieved by a mechanism not explained by the substrate specificity of MurC. Our results showing YfiH as an L-Ser-specific UDP-MurNAc-monopeptide amidase therefore uncover a bacterial control mechanism to ensure only the presence of the precursor UDP-MurNAc-L-Ala in the peptide assembly pathways by removing unwanted L-Ser containing PG-precursors made by the MurC reactions. It is therefore likely that bacteria rely on specific YfiH-like hydrolases to maintain a PG with specific stem peptides. Finally, it remains to be seen whether bacterial cells can regulate expression of YfiH in response to environmental stimuli to alter the composition of the first amino acid of the PG stem peptide, thereby evading the attack of muralytic enzymes to gain survival advantage.

**Figure 5.**
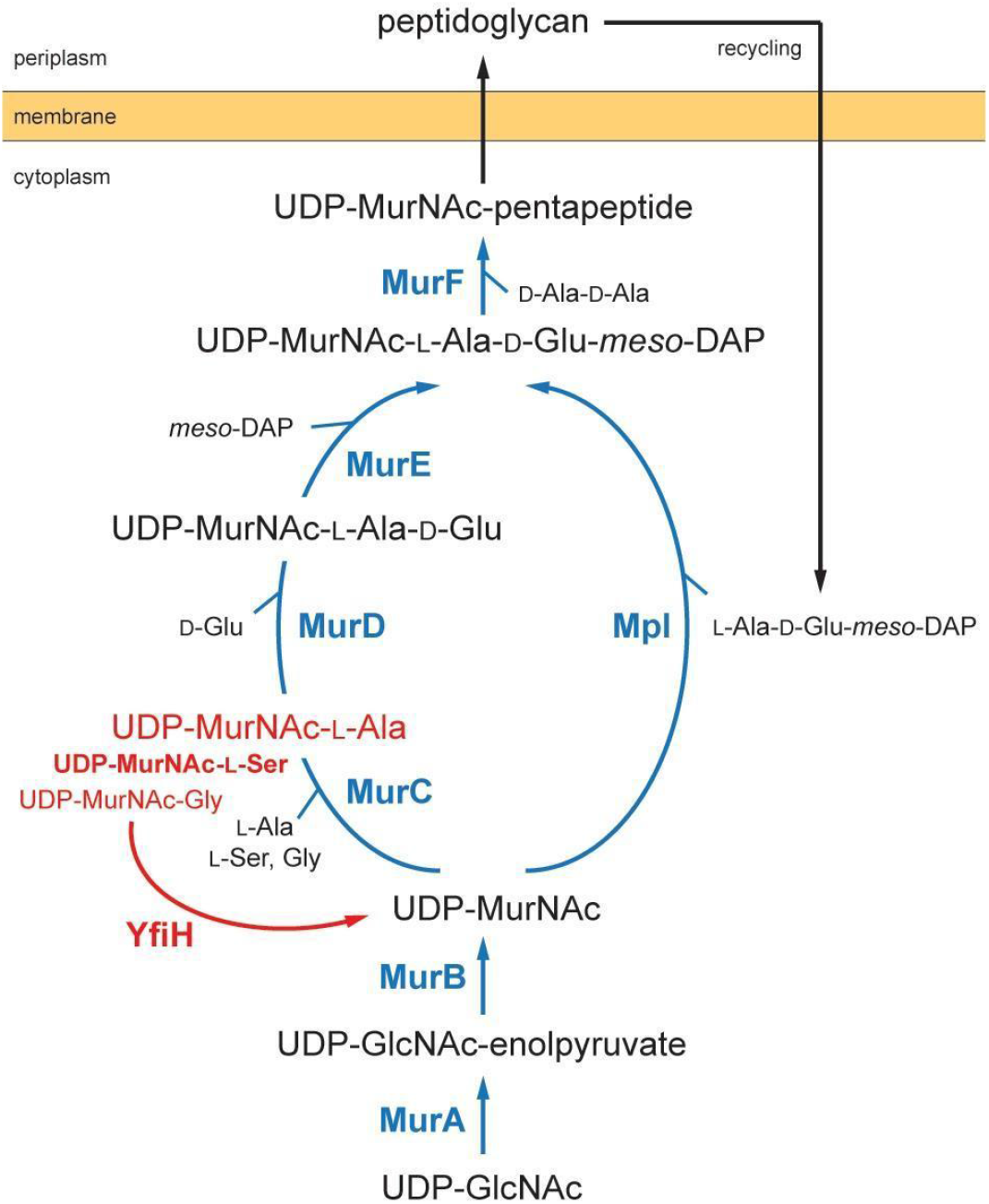
Involvement of YfiH in the cytoplasmic steps of peptidoglycan biosynthesis and recycling. The reactions catalyzed by the ATP-dependent Mur/Mpl ligases are colored in blue. The reactions catalyzed by YfiH are highlighted in red.

YfiH belongs to a family of proteins containing the domain of unknown function (DUF) 152, classified in the Pfam database. Our structural and functional analysis of *E. coli* YfiH uncovers a binding groove for the PG precursor UDP-MurNAc-monopeptide. Interestingly, YfiH also cleaves MurNAc-L-Ser (Fig. S2A). This result suggests that the conserved L-shaped binding groove formed in the structure of DUF152 proteins may bind extracytoplasmic PG-related fragments, which may be mechanistically important for the function of some homologues found in higher eukaryotic organisms. One possibility is that some of the DUF152 proteins may have lost enzymatic activity but evolved to retain specific ligand-binding activity as seen in the peptidoglycan recognition proteins (PGRPs), which play sensor roles in innate immune recognition (Dziarski 2004). FAMIN, a DUF152 protein, is shown to be overexpressed in macrophages under muramyl dipeptide (MDP) treatment and associated with NOD2-induced intracellular microbial clearance (Lahiri et al. 2017). Based on a homology model of FAMIN-CTD predicted by AlphaFold, we have noticed that in the conserved binding groove, the specificity determinant residue is a threonine (Thr74) in lieu of glutamine; moreover, the location of the arginine likely interacting with the carboxylic group of the amino-acid moiety of the PG precursor is different from that of Arg235 of YfiH, which may be essential for catalytic activity (Figs. 2 and S3). The misplacement of the substrate carboxyl group-coordinating basic residue and the change of specificity determinant residue in FAMIN-CTD suggest that it may function as a sensor recognizing UDP-MurNAc-L-Ala or MurNAc-L-Ala. Future studies will be needed to characterize the PG-recognition activities of these DUF152 proteins and their biological roles.

## Materials and Methods

### Cloning and mutagenesis

All plasmids in this study were subcloned into a pET21a(+) vector with C-terminal 6xHis-tag. YfiH-C107A, YfiH-Q69A, and YfiH-Q69M were generated using the wild-type YfiH (UniportKB:P33644) plasmid as the template by PCR-based site-directed mutagenesis. Gene encoding MurC (UniProtKB: P17952) and MurD (UniProtKB:P14900) were cloned from *E. coli BL21(DE3)* competent cells by PCR. All constructs in this study were sequenced prior to use by DNA Sequencing Core Facility of Academia Sinica (AS-CFII-108-115).

### Protein expression and purification

Plasmid was transformed into ECOS™ *E. coli BL21(DE3)* cells (Yeastern Biotech). Cells were cultured to an optical density at 600 nm of 0.6–0.8 and induced with 1 mM isopropyl β-D-thiogalactopyranoside at 20 °C for 18 h. Cell pellets were resuspended in lysis buffer containing 50 mM Tris-HCl, pH 8.0, 500 mM NaCl with protease inhibitor cocktail (Roche) and then ruptured by French press (Avestin). After centrifugation at 35,000 × g at 4 °C for 45 min, the supernatant was applied to a nickel-nitrilotriacetic acid agarose (Qiagen) column and washed with 20 mM imidazole for twice. The protein fraction eluted with 250 mM imidazole was purified further by MonoQ 5/50 GL column (GE Healthcare) at pH 8.0, and Superdex 200 10/300 GL column (GE Healthcare) was equilibrated in 20 mM Tris-HCl, pH 8.0, 100 mM NaCl and 1 mM DTT.

### Crystallization, structure determination, and refinement

The crystallization of YfiH-C107A was performed by using sitting-drop vapor diffusion method at 22 °C, in which 1 μl 15 mg/mL protein solution was mixed with 1 μl reservoir solution consisting of 2.0 M ammonium sulfate and 100 mM sodium acetate at pH 5.4. The crystals of YfiH-C107A bound to UDP-MurNAc-Ser were prepared by washing the YfiH-C107A crystals, which contain bound endogenous UDP-MurNAc, in 10 μl of the mother liquid twice, followed by transferring the crystals to a 5 μl drop of freshly prepared reservoir solution consisting of 2.2 M ammonium sulfate and 100 mM sodium phosphate at pH 5.4, added with 0.25 μl of 20 mM of UDP-MurNAc-Ser, and incubated for 1.5 weeks; the procedure was repeated twice. The crystals were cryoprotected by a brief transfer to the mother liquid supplemented with 10-30% xylitol prior to data collection.

All diffraction data were collected on NSRRC beamline 15A1 (NSRRC Taiwan). All images were indexed, integrated, and scaled using the HKL-2000 package (Otwinowski and Minor 1997). The structures were solved by molecular replacement with the program PHASER (McCoy et al. 2007). The crystal structure of *Escherichia Coli* YfiH (PDB:1Z9T) was used as the initial search model. Structure refinement and manual modeling were implemented using the programs Refmac5 and COOT, respectively (Murshudov et al. 2011; Emsley and Cowtan 2004). Native ligands bound in crystal structures were explored using the program LigandFit in Phenix. The cut-off of minimum correlation coefficient was setted to 0.75 to avoid incorrected ligand placement. All protein model figures were generated with PyMOL (v.1.7.2; Schrödinger).

### Synthesis of UDP-MurNAc-peptide derivatives

UDP-MurNAc-Ala and UDP-MurNAc-Ser were produced by enzymatic synthesis. The processes were performed in a reaction mixture containing 20mM Tris-HCl pH8.0, 0.5mM UDP-MurNAc (Chiralix), 5mM ATP, 10mM MgCl_2_ and 0.01mM MurC enzyme, follow by adding 1mM L-alanine or L-serine as amino acid donor. The reaction was performed completely at 37°C for 18 hours then stopped the reaction by removing enzymes with Amicon® Ultra-15 Centrifugal Filter Unit (Millipore). The product was purified by reverse phase HPLC. Fraction with desired UDP-MurNAc-amino acid was collected at 262 nm and reconfirmed by ESI-MASS. The qualified fractions were lyophilized and reconstituted in de-ionized water. Synthesis of UDP-MurNAc-Ala-Glu or UDP-MurNAc-Ser-Glu were similar as described above, but with extra 2 mM D-Glutamate and 0.5mM MurD enzyme during the reaction step.

### Procedures to synthesize the MurNAc-Ser derivatives

All reactions were conducted in oven-dried glassware, under nitrogen atmosphere (Scheme S1). The reaction products were purified by using column chromatography on silica gel (Geduran Silicagel 60, 0.040-0.063 mm, from Geduran®), Buchi Puro850 automated purification machine or HPLC. Anhydrous solvents and moisture-sensitive materials were transferred by using an oven-dried syringe or cannula through a rubber septum. Organic solutions were concentrated under reduced pressure in a water bath (< 40 °C). TLC was performed on pre-coated glass plates of TLC Silica gel 60G F254 (Merck KGaA®), and was detected by UV lamp (254 nm) and/or by staining reagents that contained ceric ammonium molybdate (for general use), *p*-anisaldehyde (for sugars) or ninhydrin (for amine or amide). ^1^H NMR spectra were recorded on Bruker AVII-500 (500 MHz) spectrometers by using CD_3_OD (d_H_ = 3.31 ppm, central line of a quintet) or D_2_O (d_H_ = 4.80 ppm) as internal standards. High-resolution mass spectroscopy (HRMS) was performed on Bruker Bio-TOF III (ESI-TOF) spectrometers and is reported as mass/charge (*m/z*) ratios with percentage relative abundance.

The solvents for extraction and chromatography were of ACS grade. Anhydrous *N,N*-dimethylformamide (DMF) was purchased from Aldrich Chemical Co. in a sealed package and stored at electronic dry box. *N*-acetylmuramic acid (MurNAc), *N*-hydroxysuccinimide (NHS), *N,N*-diisopropylcarbodiimide (DIC), H-Ser-O^t^Bu,HCl salt, triethylamine (TEA), triisopropylsilane (TIS) and trifluoroacetic acid (TFA) were purchased from Bachem, BLD, TRC, BLD, J-T Baker, Aldrich and Alfa asear chemical Co., respectively. All the chemicals were directly used without further purification unless otherwise specified.

### Enzyme activity assay

In the assay, 3 nM of YfiH enzymes were incubated with 10 μM UDP-muropeptides derivatives in a reaction buffer containing 20mM Tris-HCl pH8.0 and 2mM DTT, the final volume in each assay was adjust to 50 μl then incubated in 37°C for 15 mins, except the condition of wildtype YfiH with UDP-MurNAc-Ser was incubated for 7mins due to unexpectedly high activity. All reactions were paused immediately by freeze in liquid nitrogen until HPLC detection.

Efficiency of substrate hydrolysis in each reaction was quantified by monitoring the change of initial peak areas under 262 nm by HPLC. Peak areas for analyte were proportional to the injected amount and were confirmed at initial assay establishment. Fraction elutes were further analyzed by mass spectrometry to address the identity of final reaction products in assays.

### Electrospray ionization-mass spectrometry (ESI-MS) analysis

The experiments aiming for high resolution and high mass accuracy in this study were done on a LTQ Orbitrap XL ETD mass spectrometer (Thermo Fisher Scientific) equipped with standard ESI ion source. 5μl of samples were flow injected at a rate of 50 μl min^−1^. in 80% ACN/H2O 0.1%FA by the ACQUITY UPLC system from Waters (Waters). Full-scan MS condition was mass range m/z 200-2000, resolution 60,000 at m/z 400. Electrospray voltage was maintained at 4 kV and capillary temperature was set at 275 °C.

### High performance liquid chromatography (HPLC)

The experiments of reverse phase HPLC in this study were done on a Waters Alliance 2695 separation module (Waters) with Hypersil BDS C18 Column (Thermo Fisher Scientific) and wBondapack C18 column (Waters). Samples were injected into the column with an isocratic flow of 50 mM ammonium formate pH 4.3 at 1 ml min^-1^. Quantification of the peak area was analyzed with an Empower 3 Data Chromatography Software (Waters).

## Data availability

The structural factors and coordinates have been deposited in the Protein Data Bank under the accession codes 7F3V and 7W1G for the complexes of YfiH with YfiH-UDP-MurNAc and YfiH-UDP-MurNAc-L-Ser, respectively.

## Acknowledgments

We thank Dr. Zhijay Tu of the IBC Chemistry Core for chemical synthesis of MurNAc-L-Ser, Yu-Ling Hwang of IBC Peptide Core for assistance with HPLC analysis, the GRC Mass Core Facility of Academia Sinica for mass spectrometric analysis, and the beamline support from NSRRC, a national user facility supported by the Ministry of Science and Technology (MOST), Taiwan, ROC. This work was supported by Academia Sinica and MOST (under Grant MOST108-2320-B-001-011-MY3).

## Author Contributions

C.I.C. conceived the study. M.S.L., K.Y.H., C.I.K, and S.H.L. acquired the data. M.S.L. and K.Y.H. solved the crystal structures. M.S.L., K.Y.H., and C.I.C. analyzed and interpreted the data and prepared the manuscript. C.I.C. acquired the resources and the funding.

## Competing interests

Authors declare that they have no competing interests.

## Supplemental Figures, Table, and Scheme

**Figure S1.**
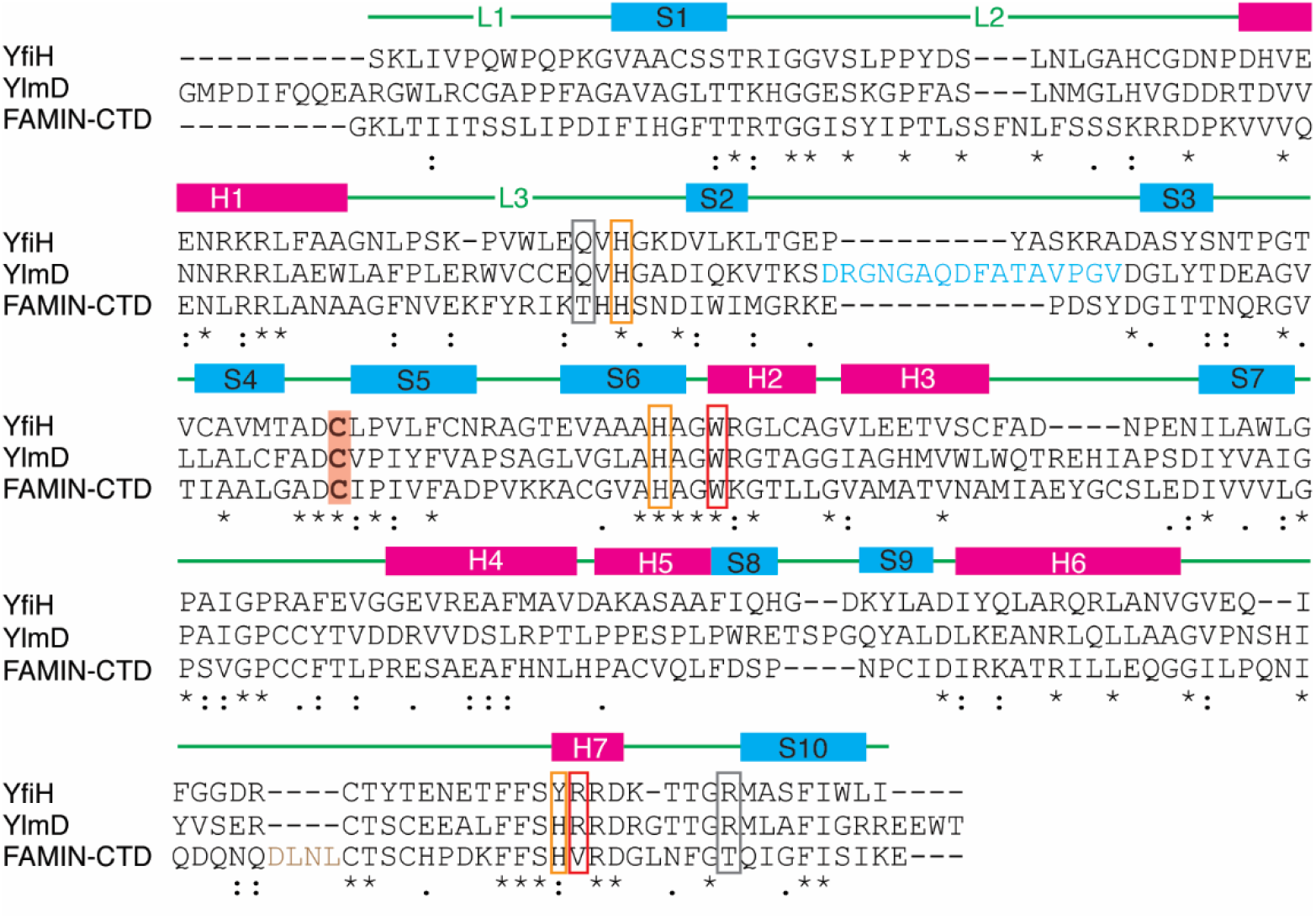
Secondary structure of YfiH and sequence alignment with YlmD and the homologous C-terminal domain (CTD) of human FAMIN/Lacc1. Secondary structure elements of YfiH are indicated above the alignment: α-helices and β-strands are illustrated by magenta and blue rectangles; loops are represented by green lines. Catalytic Cys residues are highlighted in a shaded box. Binding residues of YfiH for the moieties of UDP, MurNAc, and monopeptide of the substrate are encircled in red, orange, and gray boxes, respectively. YlmD- and FAMIN-CTD-specific insertion sequences are colored in cyan and brown, respectively. The Uniprot IDs of the aligned sequences are as follows: P33644 for *E. coli* YfiH, P84138 for *Geobacillus stearothermophilus* YlmD, and Q8IV20 for human FAMIN/Lacc1.

**Figure S2.**
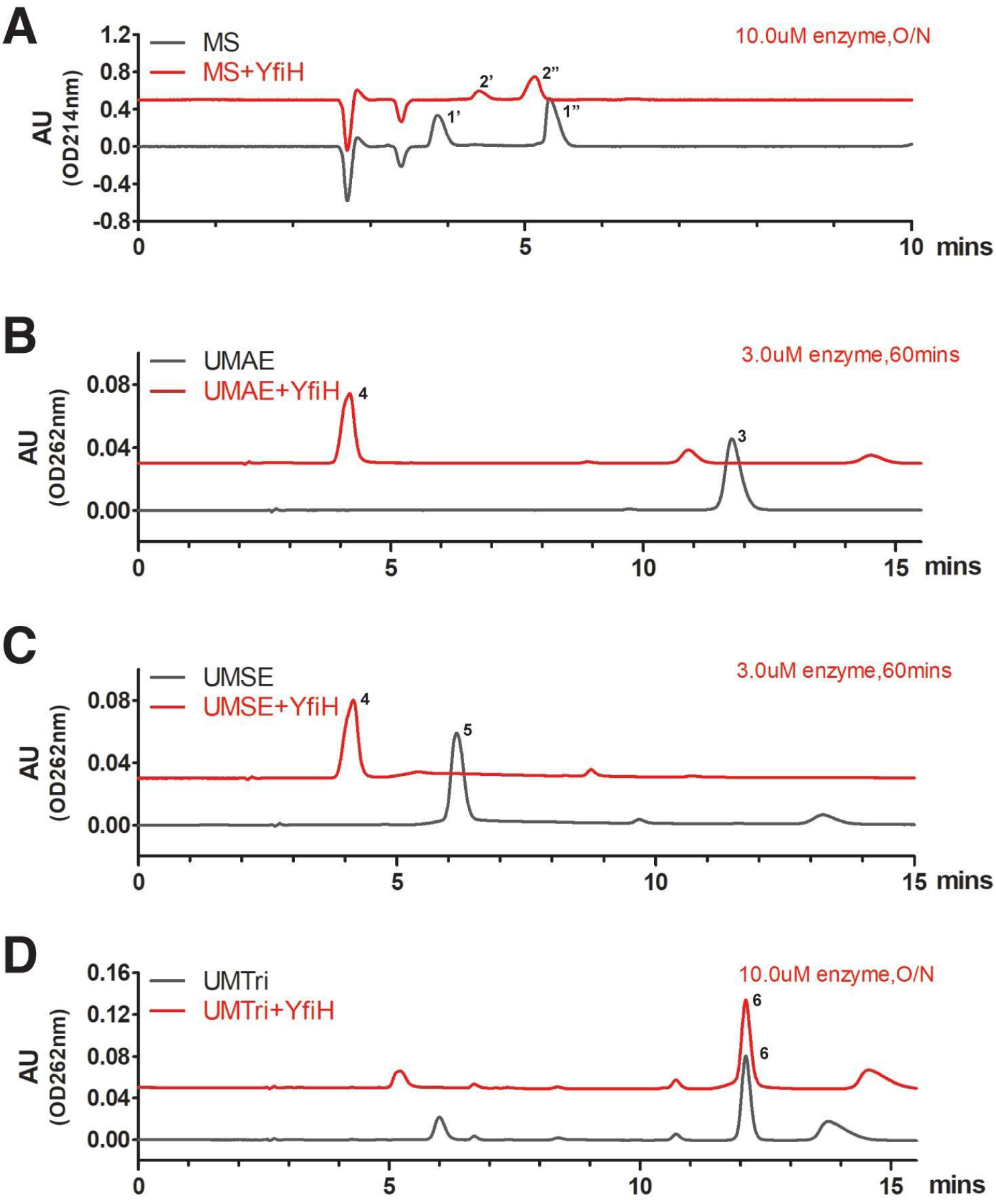
Hydrolyzing activities of YfiH on various UDP-MurNAc-peptide derivatives. A. HPLC chromatograms of MurNAc-L-Ser (MS; compounds 1’ and 1” with MurNAc in α and β anomeric forms, respectively) before and after incubation with YfiH, yielding the cleavage product MurNAc (compounds 2’ and 2”, in α and β anomeric forms, respectively). B. HPLC chromatograms of UDP-MurNAc-L-Ala-γ-D-Glu (UMAE; compound 3) before and after incubation with YfiH, yielding the cleavage product UDP-MurNAc (compound 4). C. HPLC chromatograms of UDP-MurNAc-L-Ser-γ-D-Glu (UMSE; compound 5) before and after incubation with YfiH, yielding the cleavage product UDP-MurNAc (compound 4). D. HPLC chromatograms of UDP-MurNAc-L-Ala-γ-D-Glu-*m*-DAP (UMTri; compound 6) before and after incubation with YfiH.

**Figure S3.**
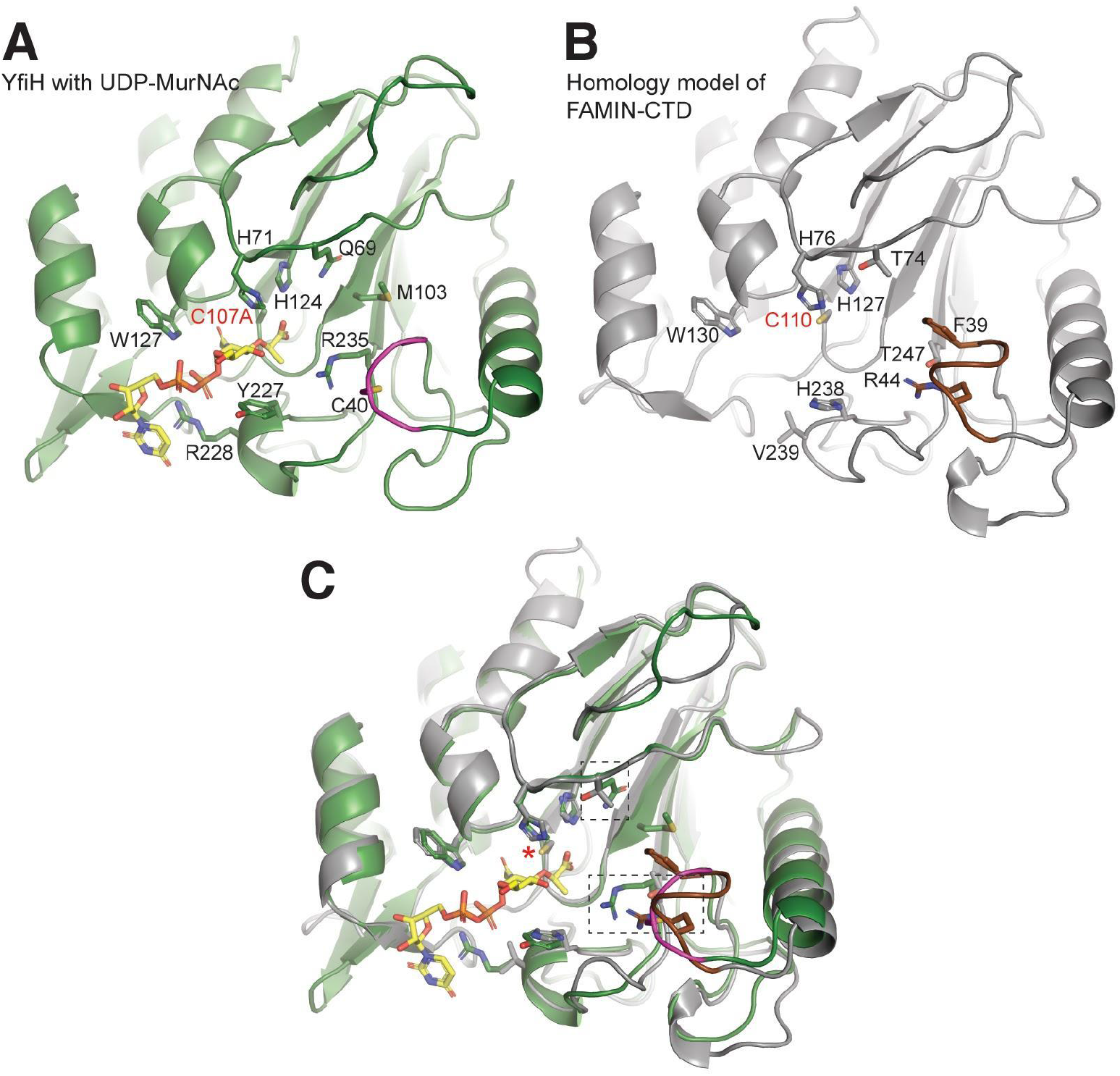
Comparison of the structure of YfiH and a homology model of FAMIN/Lacc1. A. Structure of *E. coli* YfiH bound to UDP-MurNAc. The loop region showing major structural difference with FAMIN is colored in magenta. B. A homology model of FAMIN-CTD, predicted by AlphaFold, in a similar orientation as YfiH. The FAMIN-specific insertion is highlighted in brown. C. Superimposition of the structures of YfiH and FAMIN-CTD. The bound compound and the interacting residues are shown in sticks. The catalytic Cys residue is indicated by the red label or the asterisk. The putative binding residues for the amino-acid moiety of UDP-MurNAc-monopeptide are indicated by the dash boxes.

**Scheme S1.**
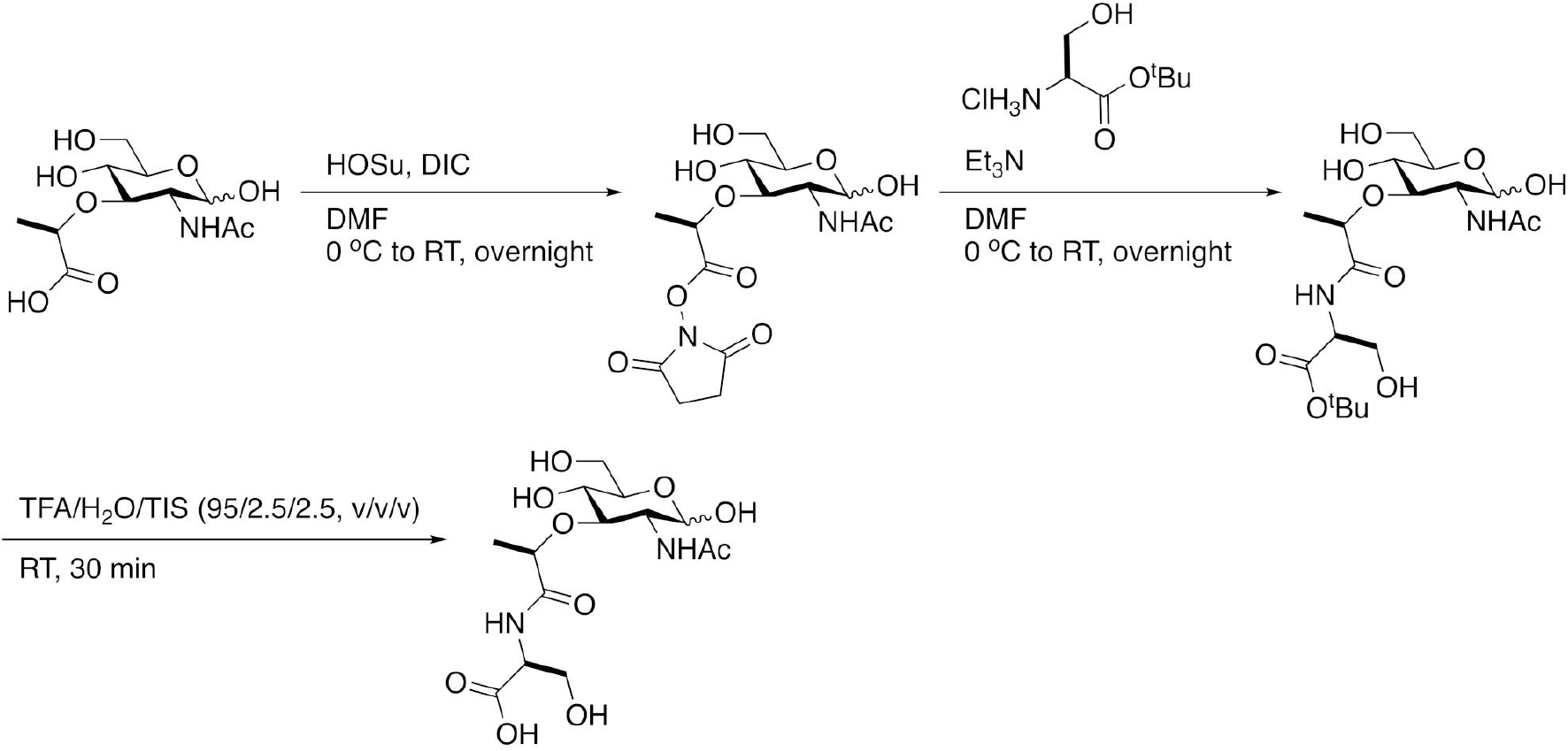
Procedures to synthesize the MurNAc-Ser derivatives. The DIC (80 μL, 0.51 mmol) was dropwise added in the solution of MurNAc (100 mg, 0.34 mmol) and NHS (47 mg, 0.41 mmol) in dry DMF (20 mL) with stirring at 0 °C under nitrogen atmosphere. The product NHS ester-activated MurNAc was further used for the next step. To the MurNAc-NHS ester (79 mg, 0.20 mmol) and H-Ser-O^t^Bu,HCl salt (80 mg, 0.40 mmol), in which the TEA (112 μL, 0.80 mmol) was dropwise added at 0 °C under nitrogen atmosphere with stirring, and further purified to obtain the desired MurNAc-Ser-OtBu (43 mg, 49% yield). The MurNAc-Ser-OtBu (6 mg, 14 μmol) then reacted with 200 μL of acid recipe TFA/H_2_O/TIS (95/2.5/2.5, v/v/v) to synthesis the target compound MurNAc-Ser-OH (0.5 mg, 10% yield).

**Table S1.** Crystallographic data collection and refinement statistics.

|  | **YfiH-C107A-UM** | **YfiH-C107A-UMS** |
| --- | --- | --- |
| **Data collection** |  |  |
| Wavelength (Å) | 1.00 | 1.00 |
| Space group | P21 21 21 | P21 21 21 |
| Cell dimensions |  |  |
| *a*, *b*, *c* (Å) | 69.95, 98.83, 136.24 | 70.56, 99.24, 136.93 |
| $\alpha$, $\beta$, $\gamma$ (°) | 90, 90, 90 | 90, 90, 90 |
| Resolution (Å) | 50-1.47 (1.52-1.47)<sup>a</sup> | 27.44-1.86 (1.927 -1.86) |
| *I*/σ*I* | 40.08 (2.4) | 10.46 (3.20) |
| *R*<sub>merge</sub> | 0.033 (0.529) | 0.130 (0.755) |
| *R*<sub>pim</sub><sup>b</sup> | 0.017 (0.27) | 0.059 (0.311) |
| CC<sub>1/2</sub><sup>c</sup> | 0.965 (0.854) | 0.963 (0.852) |
| Completeness (%) | 97.3 (94.0) | 92.71 (90.25) |
| Redundancy | 6.9 (6.7) | 5.1 (5.6) |
| **Refinement** |  |  |
| Resolution (Å) | 34.08-1.47 (1.51-1.47) | 27.44 -1.86 (1.93 - 1.86) |
| No. of reflections | 141889 (7130) | 75418 (7224) |
| Reflections used for *R*<sub>free</sub> | 7531 (356) | 3695 (354) |
| *R*<sub>work</sub> | 0.185 (0.255) | 0.2199 (0.2872) |
| *R*<sub>free</sub> | 0.214 (0.28) | 0.2610 (0.2987) |
| No. atoms |  |  |
| Protein | 7297 | 7435 |
| Water | 838 | 444 |
| B-factors (Å<sup>2</sup>) |  |  |
| Protein | 16.88 | 33.98 |
| Water | 26.32 | 36.07 |
| R.m.s. deviations |  |  |
| Bond lengths (Å) | 0.013 | 0.008 |
| Bond angles (°) | 1.857 | 1.574 |
| Validation |  |  |
| Clash score | 4 | 5.07 |
| Rotamer outliers (%) | 1.74 | 3.01 |
| Ramachandran plot |  |  |
| Favored/Allowed/Disallowed (%) | 96.2/3.3/0.5 | 95.52/4.06/0.42 |

<sup>a</sup> Highest resolution shell is shown in parenthesis.
<sup>b</sup> *R*<sub>pim</sub> is the precision-indicating merging *R*, which describes the accuracy of the averaged measurement (Weiss 2001).
<sup>c</sup>CC<sup>1/2</sup> is the correlation coefficient between two random half data sets (Karplus and Diederichs 2012).

## References

Barreteau H, Kovač A, Boniface A, Sova M, Gobec S, Blanot D. 2008. Cytoplasmic steps of peptidoglycan biosynthesis. FEMS Microbiol Rev 32: 168–207.

Cabeen MT, Jacobs-Wagner C. 2005. Bacterial cell shape. Nat Rev Microbiol 3: 601–610.

Cader MZ, de Almeida Rodrigues RP, West JA, Sewell GW, Md-Ibrahim MN, Reikine S, Sirago G, Unger LW, Iglesias-Romero AB, Ramshorn K, et al. 2020. FAMIN Is a Multifunctional Purine Enzyme Enabling the Purine Nucleotide Cycle. Cell 180: 815.

Dziarski R. 2004. Peptidoglycan recognition proteins (PGRPs). Mol Immunol 40: 877–886.

Emanuele JJ Jr, Jin H, Jacobson BL, Chang CY, Einspahr HM, Villafranca JJ. 1996. Kinetic and crystallographic studies of Escherichia coli UDP-N-acetylmuramate:L-alanine ligase. Protein Sci 5: 2566–2574.

Emsley P, Cowtan K. 2004. Coot: model-building tools for molecular graphics. Acta Crystallogr D Biol Crystallogr 60: 2126–2132.

Karplus PA, Diederichs K. 2012. Linking crystallographic model and data quality. Science 336: 1030–1033.

Kim Y, Maltseva N, Dementieva I, Collart F, Holzle D, Joachimiak A. 2006. Crystal structure of hypothetical protein YfiH from Shigella flexneri at 2 A resolution. Proteins 63: 1097–1101.

Kohlrausch U, Höltje JV. 1991. One-step purification procedure for UDP-N-acetylmuramyl-peptide murein precursors from Bacillus cereus. FEMS Microbiol Lett 62: 253–257.

Lahiri A, Hedl M, Yan J, Abraham C. 2017. Human LACC1 increases innate receptor-induced responses and a LACC1 disease-risk variant modulates these outcomes. Nat Commun 8: 15614.

Liger D, Masson A, Blanot D, van Heijenoort J, Parquet C. 1995. Over-production, purification and properties of the uridine-diphosphate-N-acetylmuramate:L-alanine ligase from Escherichia coli. Eur J Biochem 230: 80–87.

Mahapatra S, Crick DC, Brennan PJ. 2000. Comparison of the UDP-N-acetylmuramate:L-alanine ligase enzymes from Mycobacterium tuberculosis and Mycobacterium leprae. J Bacteriol 182: 6827–6830.

McCoy AJ, Grosse-Kunstleve RW, Adams PD, Winn MD, Storoni LC, Read RJ. 2007. Phaser crystallographic software. J Appl Crystallogr 40: 658–674.

Murshudov GN, Skubák P, Lebedev AA, Pannu NS, Steiner RA, Nicholls RA, Winn MD, Long F, Vagin AA. 2011. REFMAC5 for the refinement of macromolecular crystal structures. Acta Crystallogr D Biol Crystallogr 67: 355–367.

Otwinowski Z, Minor W. 1997. [20] Processing of X-ray diffraction data collected in oscillation mode. Methods Enzymol 276: 307–326.

Parveen S, Reddy M. 2017. Identification of YfiH (PgeF) as a factor contributing to the maintenance of bacterial peptidoglycan composition. Mol Microbiol 105: 705–720.

Schleifer KH, Kandler O. 1972. Peptidoglycan types of bacterial cell walls and their taxonomic implications. Bacteriol Rev 36: 407–477.

Vollmer W, Blanot D, de Pedro MA. 2008. Peptidoglycan structure and architecture. FEMS Microbiol Rev 32: 149–167.

Weiss MS. 2001. Global indicators of X-ray data quality. Journal of Applied Crystallography 34: 130–135. http://dx.doi.org/10.1107/s0021889800018227.

